## Supplemental figures for "Negative Regulation of Autophagy by UBA6-BIRC6–Mediated Ubiquitination of LC3"

### Supplementary information

#### Supplementary antibodies

Antibodies to the following antigens (supplier and catalog information in parenthesis) were used in the supplementary experiments: LC3A (Cell Signaling Technology, 4599), GABARAP (Cell Signaling Technology, 13733), GABARAPL1 (Cell Signaling Technology, 26632), ATG7 (Cell Signaling Technology, 8558), ATG3 (Cell Signaling Technology, 3415), ATG16 (Cell Signaling Technology, 8089), ATG5 (Cell Signaling Technology, 12994), ATG12 (Cell Signaling Technology, 4180), WIPI2 (Bio-Rad, MCA5780), p-BECN1 (Abbiotec, 254515), BECN1 (Cell Signaling Technology, 3495), ATG14 (MBL International, PD026), p-ATG13 (Rockland Immunochemicals, 600-401-C49S), ATG13 (Cell Signaling Technology, 13468), p-ULK1 (Cell Signaling Technology, 12753), ULK1 (Cell Signaling Technology, 8054), p-S6K (Cell Signaling Technology, 9234) , S6K (Cell Signaling Technology, 2708), p-4EBP (Cell Signaling Technology, 2855), 4EBP (Cell Signaling Technology, 9452), p-TSC2 (Cell Signaling Technology, 3617), TSC2 (Cell Signaling Technology, 4308), p-AKT (Cell Signaling Technology, 13038), AKT (Cell Signaling Technology, 4691), ATG9A (Cell Signaling Technology, 9730), TGN46 (Bio-Rad, AHP500GT), Calnexin (EMD Millipore, MAB3126), GM130 (BD Biosciences, 610823), LAMTOR4 (Cell Signaling Technology, 12284), Rab5 (Cell Signaling Technology, 2143), APPL1 (Santa Cruz Biotechnology, sc-271901), EEA1 (Cell Signaling Technology, 3288), Alexa Fluor 647 Phalloidin (ThermoFisher Scientific, A22287).

**Figure S1 (related to Figure 1).**

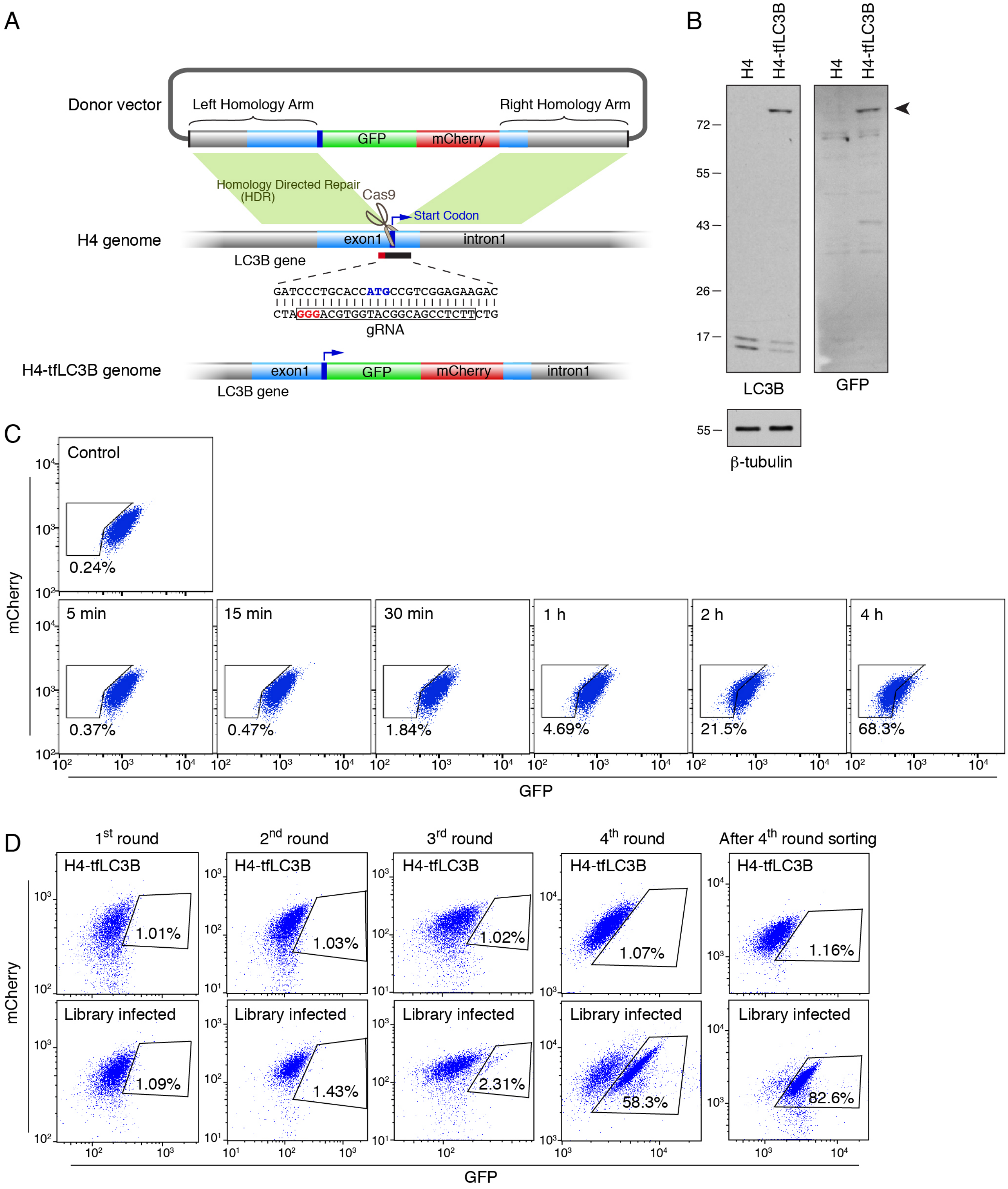

**Figure S1 (related to Figure 1). Generation, analysis and mutant selection of H4 cells expressing endogenously-tagged GFP-mCherry-LC3B.**

**(A)** Schematic representation of the generation of GFP-mCherry-tagged LC3B in H4 neuroglioma cells (H4-tfLC3B) by CRISPR/Cas9-mediated genome editing. A donor vector for HDR (homology-directed repair) was constructed by inserting a DNA fragment encoding the left homology arm, GFP-mCherry, and the right homology arm into pCI-neo vector (E1841, Promega). A double-strain break on the genomic DNA was generated by Cas9 nuclease guided by gRNA targeting the first exon of the LC3B-encoding gene. The break was repaired by HDR using donor vector as a template. **(B)** Immunoblotting of H4 and H4-tfLC3B cells with antibodies to LC3B, GFP and β-tubulin (loading control). The arrowhead indicates the position of the GFP-mCherry-LC3B protein. The positions of molecular mass markers (in kDa) are indicated on the left. **(C)** H4-tfLC3B cells were deprived of amino acids and serum for the indicated periods and analyzed by FACS. Notice how the GFP signal declined over time, reflecting the activation of autophagy during starvation. **(D)** H4-tfLC3B cells were infected with genome-wide CRISPR/Cas9 KO lentiviral pool, and cells with increased GFP signal were collected by cell sorting. Sorted cells were propagated for the next round of sorting. The naïve H4-tfLC3B cells were used to determine gating in each sorting. Notice how the population of library-infected GFP-positive cells was enriched gradually after each sorting.

**Figure S2 (related to Figure 2).**

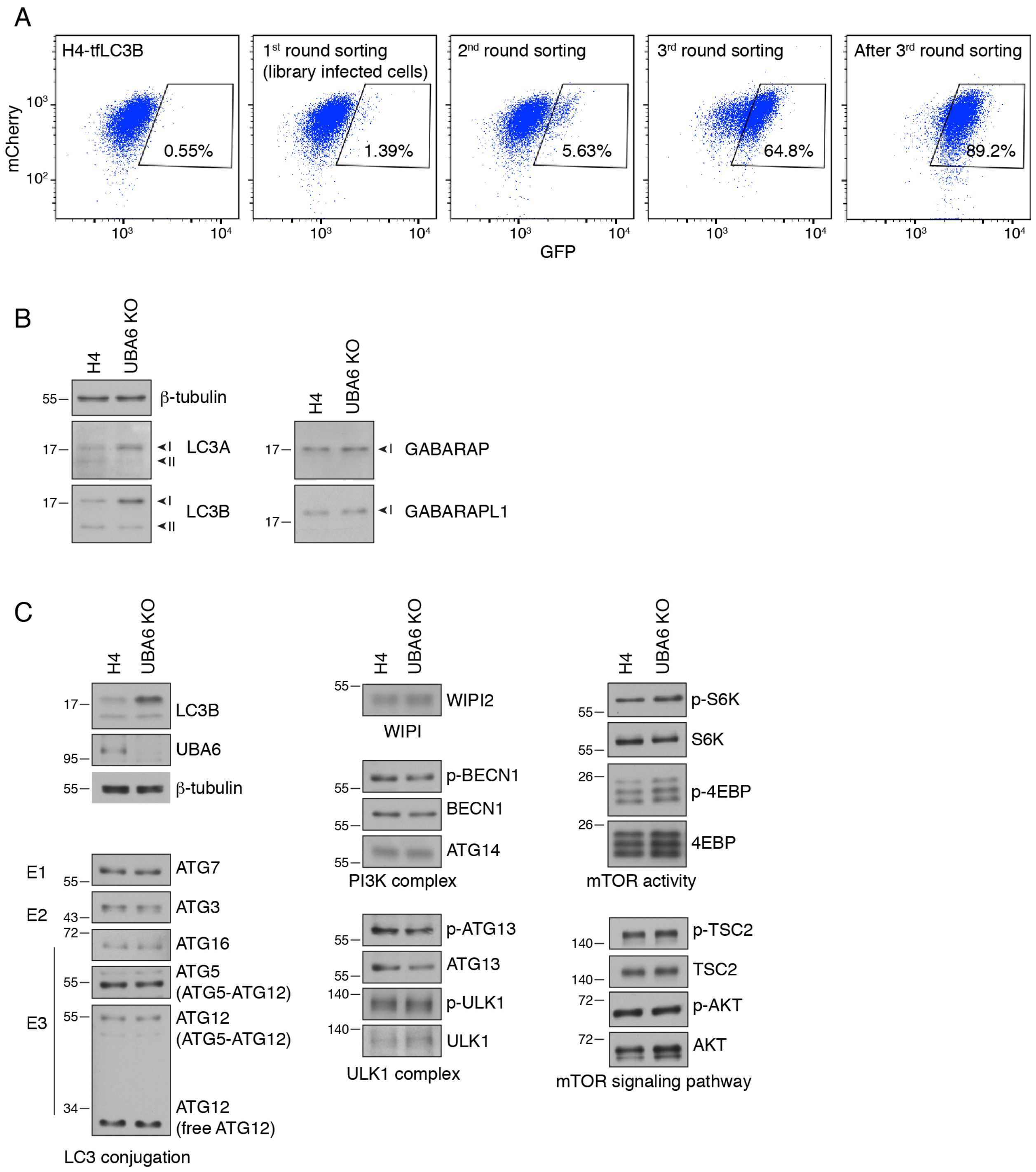

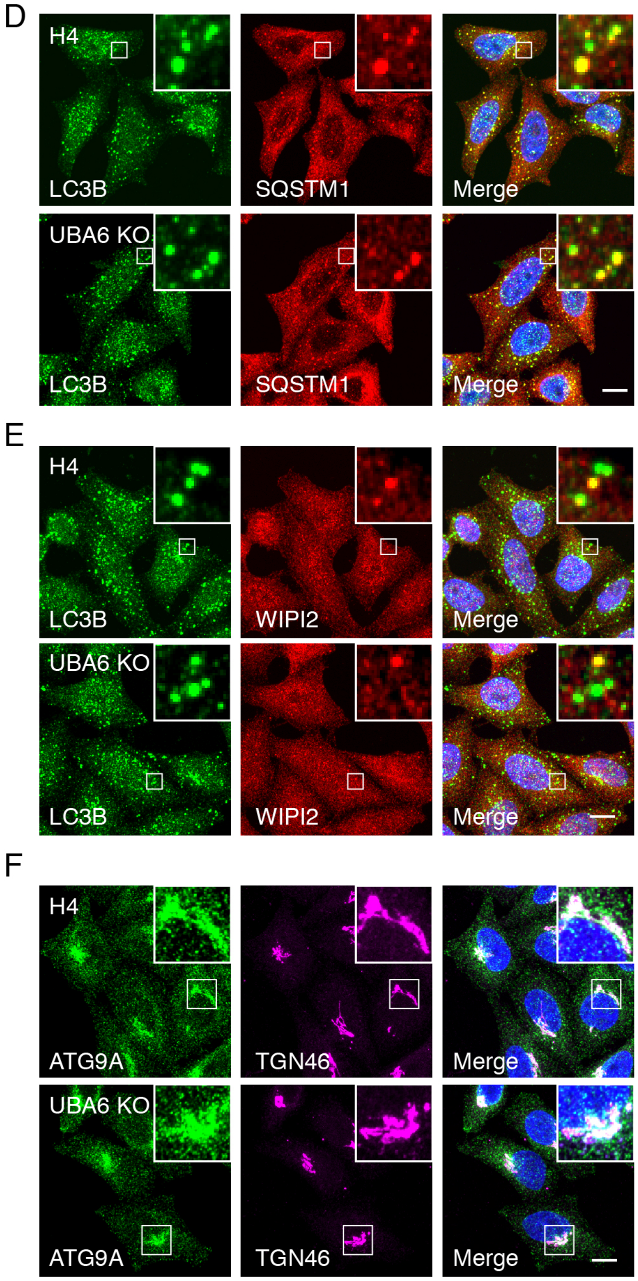
D
H4
LC3B
SQSTM1
Merge
UBA6 KO
LC3B
SQSTM1
Merge
E
H4
LC3B
WIPI2
Merge
UBA6 KO
LC3B
WIPI2
Merge
F
H4
ATG9A
TGN46
Merge
UBA6 KO
ATG9A
TGN46
Merge

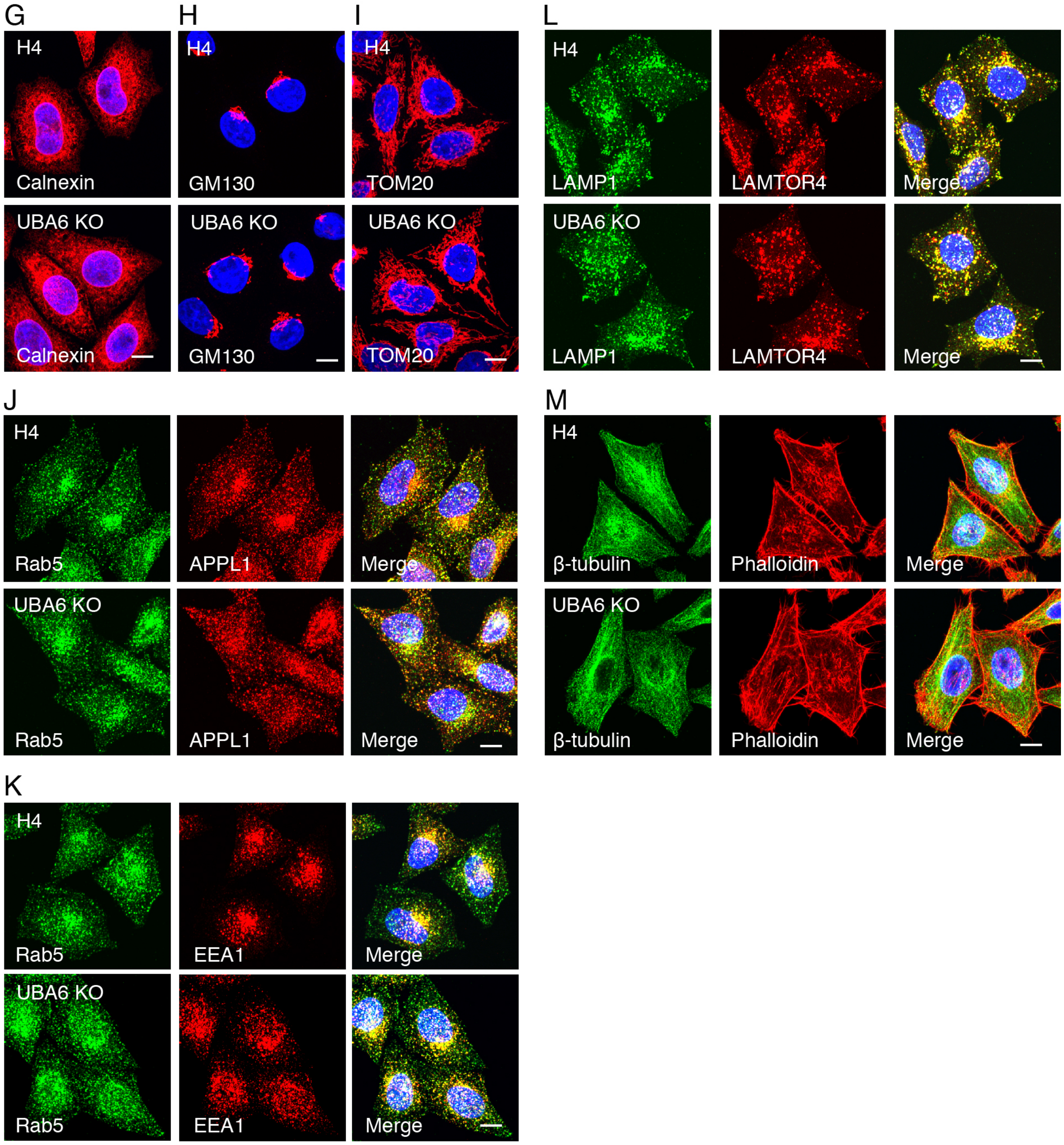
G
H
I
L
H4
Calnexin
GM130
TOM20
UBA6 KO
LAMP1
LAMTOR4
Merge
J
M
Rab5
APPL1
β-tubulin
Phalloidin
K
EEA1

**Figure S2 (related to Figure 2). Secondary screen for autophagy mutants, and expression of autophagy proteins and distribution of organellar proteins in UBA6-KO cells.**

**(A)** H4-tfLC3B cells were infected with a secondary lentiviral library targeting the top 432 genes in the primary screen. The cells with increased GFP signal were collected and enriched by 3 rounds of sorting and propagation. **(B)** WT and UBA6-KO H4 cells were lysed in 1xLDS sample buffer and analyzed by immunoblotting for Atg8 family proteins. Notice how LC3A-I and LC3B-I were increased in the KO cells. Levels of GABARAP-I and GABARAPL1-I were not elevated in UBA6-KO cells. LC3C and GABARAPL2 were not tested because of lack of good antibodies. **(C)** WT and UBA6-KO H4 cells were subjected to immunoblotting to determine the total levels or phosphorylation status of proteins involved in autophagy. Notice that none of the components of the LC3 conjugation system, WIPI2, PI3K complex, ULK1 complex and mTORC1 signaling pathway were significantly altered in UBA6-KO cells. In B and C, the positions of molecular mass markers (in kDa) are indicated on the left. **(D)** Confocal microscopy of WT and UBA6-KO H4 cells stained with antibodies to LC3B and SQSTM1. Punctate structures and colocalization of LC3B with SQSTM1 were observed in WT as well as UBA6-KO cells. Scale bar: 10 μm. **(E)** Confocal microscopy showing colocalization of LC3B and WIPI2 in WT and UBA6-KO H4 cells. Nascent phagophores were positive for both proteins, while mature autophagosomes were only positive for LC3B. Scale bar: 10 μm. **(F)** Confocal microscopy showing the distribution of ATG9A and TGN46 in WT and UBA6-KO H4 cells. In both cell lines, the majority of ATG9A localized to *trans*-Golgi network (TGN) and peripheral vesicles. Scale bar: 10 μm. **(G-M)** Confocal microscopy of WT and UBA6-KO cells stained with antibodies to calnexin (G) (endoplasmic reticulum), GM130 (H) (Golgi apparatus), TOM20 (I) (mitochondria), Rab5, APPL1, EEA1 (J, K) (early endosomes), LAMP1, LAMTOR4 (L) (late endosomes and lysosomes), β-tubulin and actin (M) (cytoskeleton). Scale bars: 10 μm. No obvious differences in staining for these markers were observed in UBA6-KO cells relative to WT cells.

**Figure S3 (related to Figure 4).**

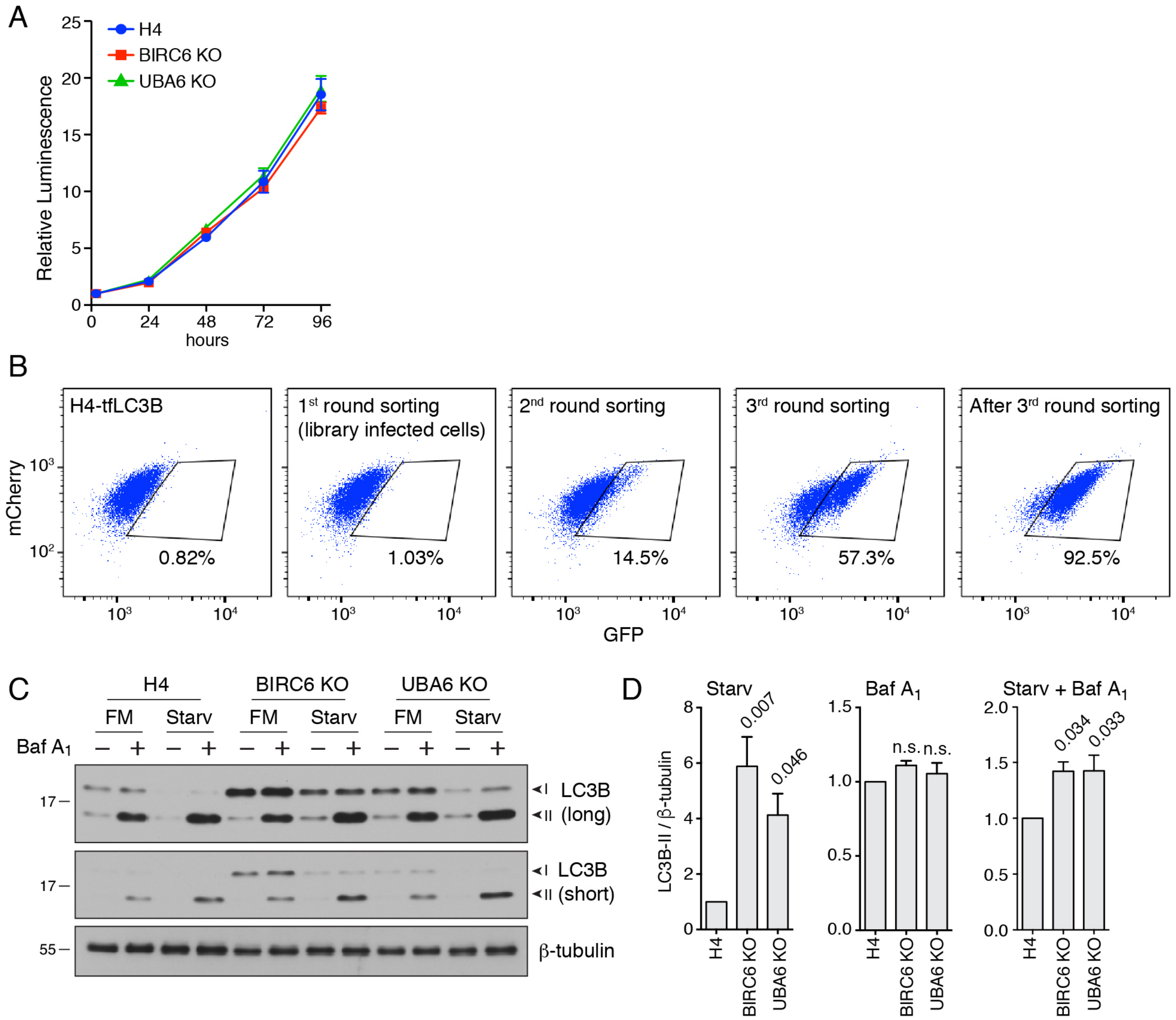


**Figure S3 (related to Figure 4). Proliferation rate of UBA6-KO and BIRC6-KO cells, screening of a ubiquitination library, and increased autophagic flux in UBA6-KO and BIRC6-KO cells.**

**(A)** Four-thousand WT, UBA6-KO and BIRC6-KO H4 cells were seeded into each well of a 96-well plate in quadruplicate for each time point in regular culture medium. After 2, 24, 48, 72 and 96 h, the number of cells per well was measured using a CellTiter-Glo 2.0 Cell Viability Assay (G9242, Promega) according to the manufacturer's instructions. The luminescence from each well represented the number of cells. The luminescence of WT H4 cells at 2 h was set as 1. Values are the mean ± SEM from 3 independent experiments. The results show that WT, UBA6-KO and BIRC6-KO H4 cells proliferated at the same rates. **(B)** H4-tfLC3B cells were infected with the ubiquitination lentiviral library targeting 661 major ubiquitination related genes. Cells with increased GFP signal were collected and propagated for several rounds of sorting. After 3 rounds, GFP-positive cells were enriched from 1.03% to 92.5%. **(C)** WT, UBA6-KO and BIRC6-KO H4 cells were incubated with bafilomycin $A_1$, nutrient-depletion medium, or a combination of both, for 2n. Cells were then lysed in 1xLDS sample buffer for immunoblotting. FM: fed medium; Starv: starvation medium lacking amino acids and serum. The positions of molecular mass markers (in kDa) are indicated on the left. **(D)** Bar graphs show the normalized LC3B levels for each condition. The indicated *p*-values were calculated using a one-way ANOVA with Dunnett's multiple comparisons test. More LC3B-II was present in UBA6-KO and BIRC6-KO cells after starvation, indicative of increased autophagy.

**Figure S4 (related to Figure 6).**

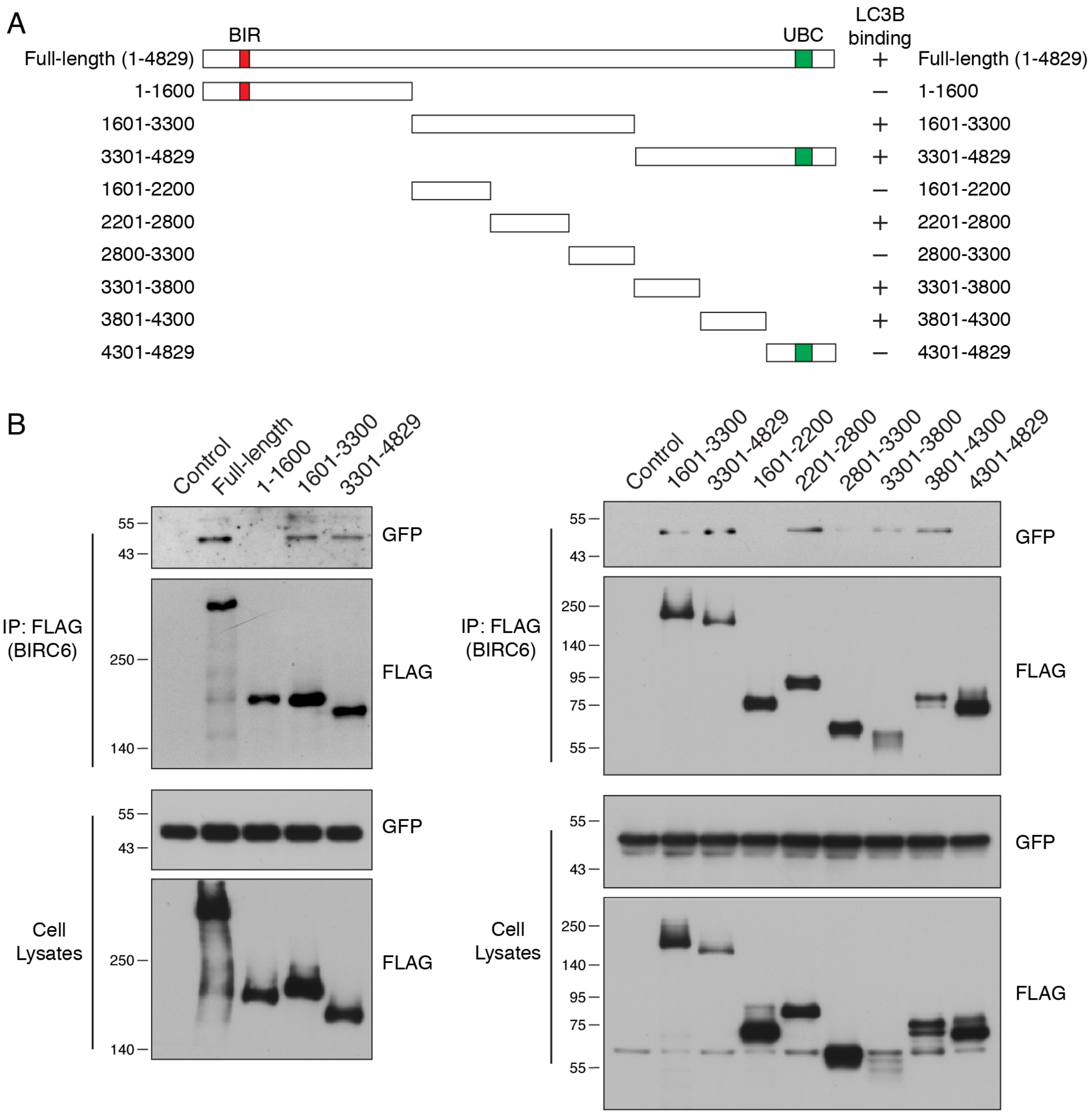


**Figure S4 (related to Figure 6). Mapping regions of BIRC6 that are required for binding to LC3B.**

**(A)** Schematic representation of full-length and truncated BIRC6. The BIR [baculoviral IAP (inhibitor of apoptosis) repeat] domain is labeled in red, and the UBC (ubiquitin-conjugating) domain is labeled in green. **(B)** HEK293T cells were co-transfected with plasmids encoding GFP-LC3B and full-length or truncated FLAG-BIRC6 deletion mutants. Immunoprecipitation was performed using antibody to the FLAG epitope. Cell lysates and immunoprecipitates were analyzed by immunoblotting with the indicated antibodies. The results suggest that the 2201-2800, 3301-3800 and 3801-4300 segments of BIRC6 participate in the interaction with LC3B. In B and C, the positions of molecular mass markers (in kDa) are indicated on the left.

**Figure S5 (related to Figure 7).**

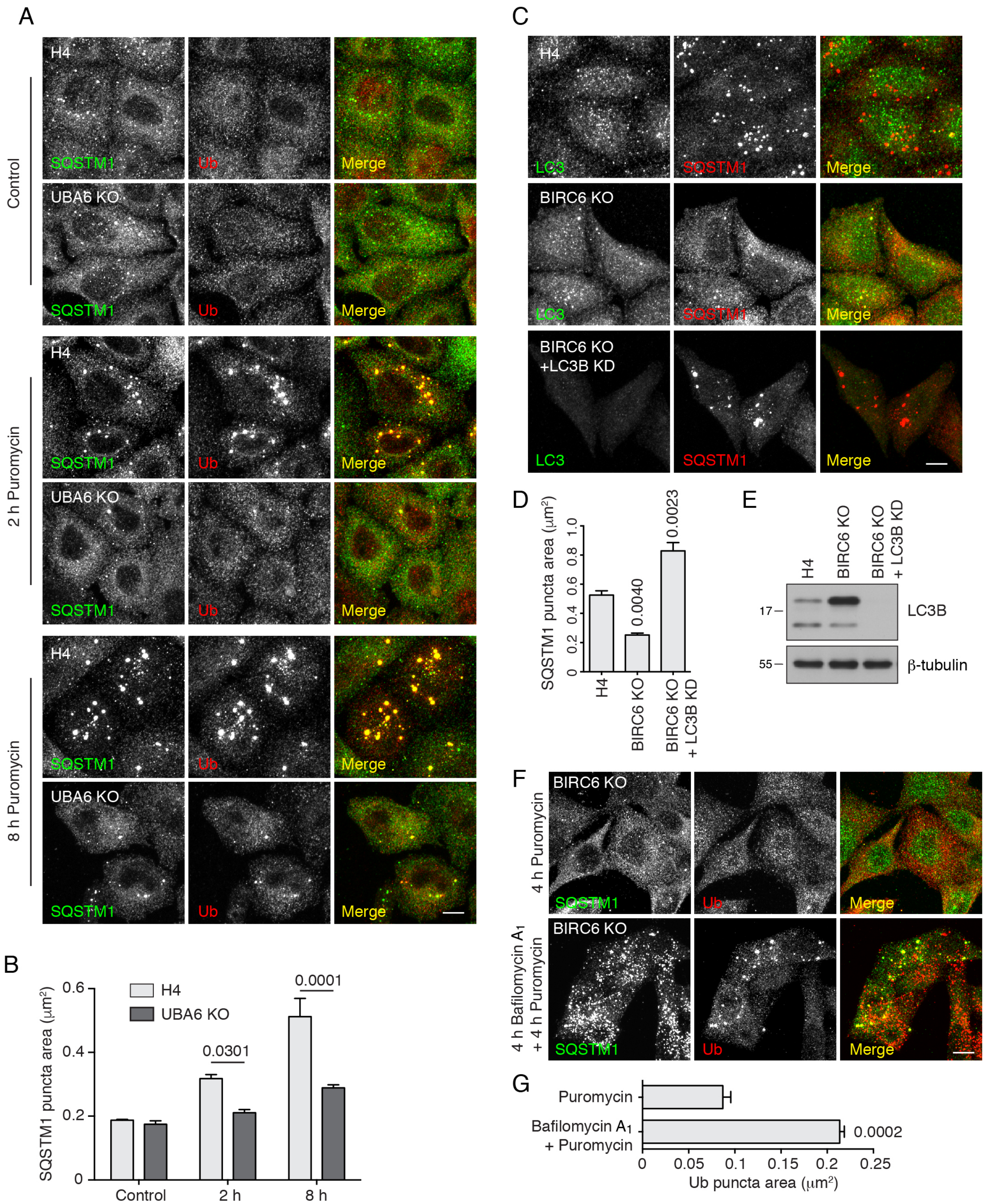

**Figure S5 (related to Figure 7). Analyses of ALIS formation in UBA6-KO and BIRC6 KO cells.**

**(A)** WT and UBA6-KO H4 cells were incubated with 5 μg/ml puromycin for 2 and 8 h. Cells were fixed and immunostained with antibodies to SQSTM1 and Ub antibodies. As shown in Figure 7, puromycin induced ALIS (aggresome-like induced structures) in WT H4 cells after a 2-h incubation. In contrast, ALIS emerged after an 8-h incubation in UBA6-KO cells. Scale bar: 10 μm. **(B)** Quantification of the area of SQSTM1-positive puncta in cells such as those shown in A. Values represent the mean ± SEM of the puncta area in 30 cells from 3 independent experiments. The indicated *p*-values were calculated using a two-way ANOVA with Tukey's multiple comparisons test. **(C)** BIRC6-KO cells were transfected with siRNA oligonucleotides targeting LC3B (6212, Cell Signaling Technology), and then incubated with 5 μg/ml puromycin for 4 h prior to immunofluorescent staining. As shown in Figure 7, puromycin did not accumulate ALIS in BIRC6-KO as compared to WT H4 cells. LC3B depletion by siRNA notably increased the accumulation of ALIS by puromycin. Scale bar: 10 μm. **(D)** Quantification of the area of SQSTM1-positive puncta in cells such as those shown in C. Values represent the mean ± SEM of the puncta area in 30 cells from 3 independent experiments. The indicated *p*-values were calculated using a one-way ANOVA with Dunnett's multiple comparisons test. **(E)** Immunoblotting of WT, BIRC6-KO and BIRC6-KO–LC3B-KD H4 cells. Notice the drastic reduction of LC3B expression in siRNA-transfected BIRC6-KO cells. **(F)** BIRC6-KO cells were pretreated with 50 nM Bafilomycin $A_1$ for 4 h prior to ALIS induction by 4-h puromycin treatment. Cells were immunostained with antibodies to SQSTM1 and Ub. Scale bar: 10 μm. **(G)** Quantification of the area of Ub-positive puncta in cells such as those shown in F. Values represent the mean ± SEM of the puncta area in 30 cells from 3 independent experiments. The indicated *p*-values were calculated using Student's *t* test. Notice that SQSTM1 puncta were prominently accumulated by bafilomycin $A_1$ pretreatment, indicating that lysosomal degradation was successfully inhibited. Furthermore, bafilomycin $A_1$-pretreated BIRC6-KO cells displayed prominent Ub puncta after 4-h puromycin incubation, indicating that the failure of BIRC6-KO cells to accumulate ALIS in the absence of bafilomycin $A_1$ is due to their higher degradative capacity relative to WT cells.

**Figure S6 (related to Figure 8).**

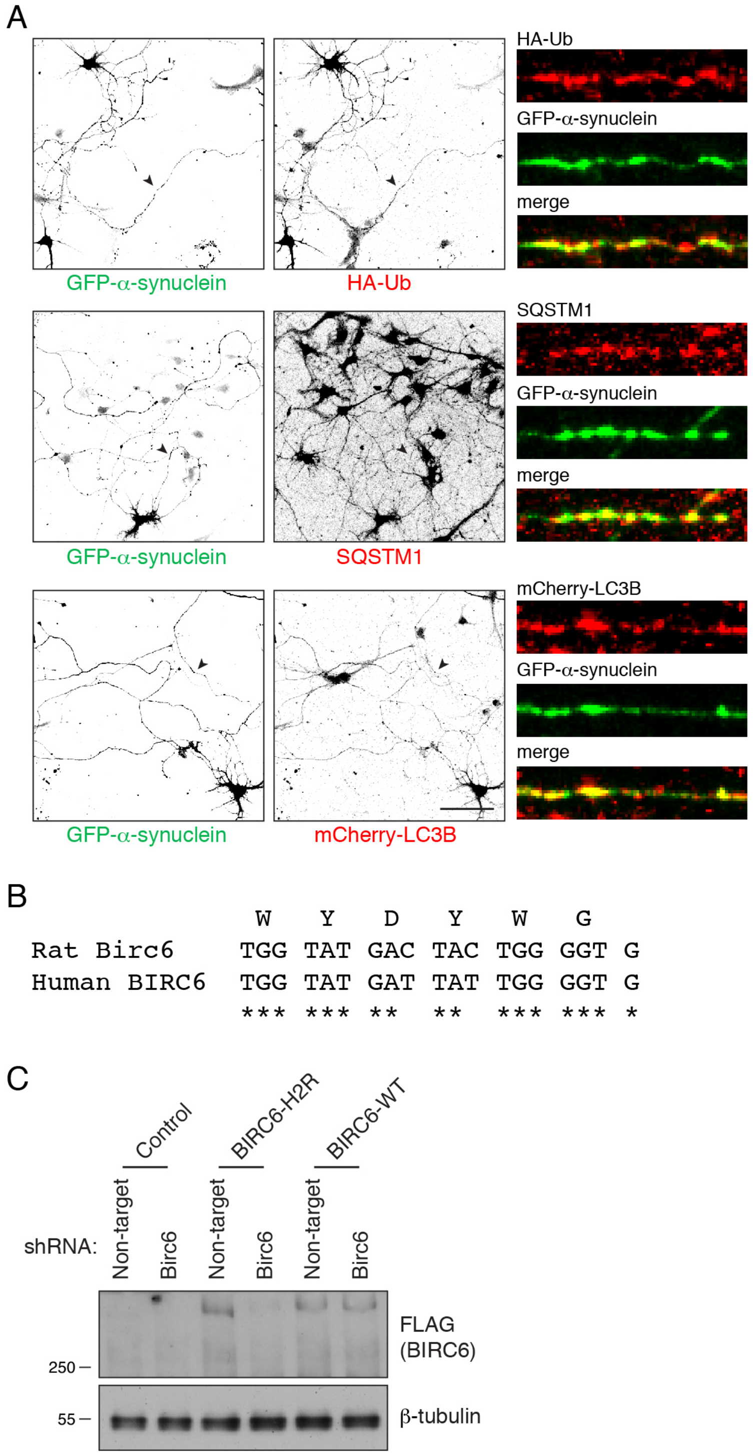

**Figure S6 (related to Figure 8). Aggregation of α-synuclein and verification of Birc6 KD in rat hippocampal neurons.**

**(A)** Rat hippocampal neurons in primary culture were cotransfected with plasmids encoding GFP-α-synuclein A53T mutant and HA-Ub or mCherry-LC3B as indicated. Transfections were performed at day-in-vitro (DIV) 3, and neurons were fixed for immunofluorescence microscopy at DIV7. Single-channel images are shown in inverted grayscale. Axons indicated by arrowheads were straightened and enlarged for better appreciation of the colocalization. Scale bar: 100 μm. Merged images show that α-synuclein aggregates colocalize with Ub, SQSTM1 and LC3B, indicating that α-synuclein aggregates are ubiquitinated and recognized for selective autophagy. **(B)** Alignment of the shRNA-target sequence of genes encoding rat Birc6 and human BIRC6. Human BIRC6 gene has a two-nucleotide mismatch as compared to the shRNA designed for rat Birc6. **(C)** BIRC6-H2R (human to rat) plasmid was generated by substituting the shRNA-targeting site in human BIRC6 with the homologous rat Birc6 sequence. H4 cells were transfected with plasmids encoding BIRC6 or BIRC6-H2R together with the plasmid encoding Birc6 shRNA. Cells were lysed in 1xLDS sample buffer and analyzed by immunoblotting with antibodies to the indicated proteins. Notice the decrease in BIRC6-H2R levels by Birc6 shRNA, verifying that the Birc6 shRNA was able to silence the expression of rat Birc6 gene.
